# Heterogeneity of Glycan Biomarker Clusters as an Indicator of Recurrence in Pancreatic Cancer

**DOI:** 10.1101/2023.01.05.522607

**Authors:** Luke Wisniewski, Samuel Braak, Zachary Klamer, ChongFeng Gao, Chanjuan Shi, Peter Allen, Brian B. Haab

## Abstract

Outcomes following tumor resection vary dramatically among patients with pancreatic cancer. A challenge in defining predictive biomarkers is to discern within the complex tumor tissue the specific subpopulations and relationships that drive recurrence. Multiplexed immunofluorescence is valuable for such studies when supplied with markers of relevant subpopulations and analysis methods to sort out the intra-tumor relationships that are informative of tumor behavior. We hypothesized that the glycan biomarkers CA19-9 and STRA, which detect separate subpopulations of cancer cells, define intra-tumoral features associated with recurrence. We probed this question using automated signal thresholding and spatial cluster analysis applied to the immunofluorescence images of the STRA and CA19-9 glycan biomarkers in whole-block tumor sections. The tumors (N = 22) displayed extreme diversity between them in the amounts of the glycans and in the levels of spatial clustering, but neither the amounts nor the clusters of the individual and combined glycans associated with recurrence. The combined glycans, however, marked divergent types of spatial clusters, alternatively only STRA, only CA19-9, or both. The co-occurrence of more than one cluster type within a tumor associated significantly with disease recurrence, in contrast to the independent occurrence of each type of cluster. In addition, intra-tumoral regions with heterogeneity in biomarker clusters spatially aligned with pathology-confirmed cancer cells, whereas regions with homogeneous biomarker clusters aligned with various non-cancer cells. Thus, the STRA and CA19-9 glycans are markers of distinct and co-occurring subpopulations of cancer cells that in combination are associated with recurrence. Furthermore, automated signal thresholding and spatial clustering provides a tool for quantifying intra-tumoral subpopulations that are informative of outcome.

## 1 Introduction

Tumor behavior varies widely among patients with pancreatic ductal adenocarcinoma (PDAC). Among patients who are diagnosed with stage I/II cancer and undergo resection plus systemic gemcitabine, 20-35% can have durable survival out to >5 years from surgery, but nearly 40% experience disease recurrence within 1 year and 10-20% have overall survival (OS) of <1 year (1,2). Treatment with modified FOLFIRINIX gives longer responses for many patients—45-55% have OS >5 years—but ~30% have recurrence times and 5-10% have OS of <1 year (2,3). The source of the differences between patients is explained in part by tumor extent: patients with lymph node involvement, positive resection margins, and high tumor size fare worse than others (4,5). Cellular and histological features such as tumor grade and perineural invasion also predict worse outcomes (4,6,7). But such conventional measures of risk stratification do not provide enough accuracy to clearly guide selection of patients who will benefit from surgery (8).

In the search for molecular markers to improve predictive accuracy, the complexity of the tumor microenvironment brings a challenge, as it has been to discern the specific features within the complexity that are driving progression. Bulk gene-expression methods uncovered subtypes of PDAC that partially explain differences between tumors (9–11). with the basal subtype showing worse outcomes that the classical subtype (6, 7). But tumors encompass a variable admixture of cells, including both non-cancer cells (10), cancer cells of more than one subtype (12,13), and cancers cells that are not easily classified by the established subtypes (14). Furthermore, cell states can interconvert based on outgrowths from progenitor populations (12), changes in microenvironment (15), or alterations in epigenetic drivers (16). To make progress in detecting predictive features considering the extreme intra-tumoral heterogeneity, methods are needed to detect specific subpopulations and the relationships between them.

Multiplexed immunofluorescence could be valuable for this purpose because it yields both a high-resolution image of the biomarker locations within the tissue and a quantification of biomarker amounts. In fact, several studies have established that the combined analysis of biomarker amounts and spatial patterns provides useful information about tumor subtyping. For example, features of immune-cell density predicted response to anti-PDL1 therapy in melanoma patients (17) and in PDAC (18), and immune cell features correlate with clinical outcome (19). But the key component for such methods to be effective is markers to detect the individual subpopulations within a tumor that have relevance to tumor behavior. Furthermore, image-analysis algorithms and software are needed to measure the levels and relationships between the markers that are informative.

Biomarkers that are promising for the detection of complementary subpopulations of cancer cells are the glycans CA19-9 and STRA (17). CA19-9 is produced by cells of the classical, epithelial, and well-differentiated type, and STRA is produced by cells of both the classical and the basal, mesenchymal and poorly differentiated types (18). CA19-9 is useful as a marker of tumor burden but not as a predictor of response. STRA, in contrast, is a marker of biological subtype and resistance to chemotherapies (18). Previous studies of primary tissue showed that tumors variously produce CA19-9, STRA, or both (17,19). The previous studies, however, did not reveal information about the whether these markers can home in the specific subpopulations of cancer cells associated with cancer progression because image analysis methods were not applied to explore the relationships among the biomarkers within the specimens.

Multiple software tools are available for analyzing multiplexed immunofluorescence images, but no particular method is established for defining cancer-cell patterns associated with outcome. The above-described studies relied on identification and annotation of histologic structures such as cell nuclei, tubules, and epithelial regions—a method known as segmentation. These methods identify cells based on color, texture, and shape based on pathologist-guided training. With proper training datasets, the methods work very well for identifying distinct cell populations. Once the cells are identified, further analyses can be performed such as nearest neighbor or density analyses. Other methods that do not require pathologist review involve the deep learning methods (20). But a potential limitation of this approach with some biomarkers is that the biomarkers do not neatly fit into cells, or that the cells sometimes have very irregular characteristics. Another concern with training is that it requires a user-defined gold standard. Such a gold standard can be hard to define, given the immense heterogeneity and diversity between tumors and the variation between pathologists in determining certain cell types (21). Variation in tissue quality also presents a problem for training (21). A complementary approach to segmentation is to analyze biomarker amounts and organization without reference to pattern recognition, but rather based on intensity threshold. The challenge with this approach is the automated setting of the intensity threshold, which is an acknowledged difficulty in the field of digital pathology (22). We previously developed an algorithm and software package that automatically determines, without user intervention, the optimal intensity threshold for an immunofluorescence image and the quantification of signal and background pixels (23). The advantage of this method is that it gives a statistically based, unbiased, consistent analysis across all images, which in turn provides a key starting point for exploring complex biomarker associations.

We hypothesized that automated threshold determination and signal identification provides a foundation for determining whether the specific subpopulations or patterns defined by the CA19-9 and STRA glycans are associated with pancreatic cancer recurrence. We tested a method that operates without user selection of locations or settings for individual specimens, as needed for an unbiased, data-guided assessment of biomarker associations.

## 2 Materials and Methods

### 2.1 Study approval and sample acquisition

The tissue samples were collected under a protocol approved by the Institutional Review Board at Duke University Medical Center. All subjects provided written, informed consent, and all methods were performed in accordance with an assurance filed with and approved by the U.S. Department of Health and Human Services. Tissue from tumor resections were formalin fixed and embedded into paraffin blocks according to standard procedures. The status, procedures, and outcomes of the patients were recorded for at least 3 years.

### 2.2 Immunostaining and fluorescence imaging

We performed immunofluorescence on 5 □m thick sections cut from formalin-fixed, paraffin-embedded blocks. Paraffin was removed from 5 μm thick FFPE sections using CitriSolv Hybrid (Decon Labs, King of Prussia, PA) containing d-limonene and isopropyl cumene, and the tissue was rehydrated through an ethanol gradient of 100%, 95%, and 70% followed by washing with PBS. Following rehydration, antigen retrieval was achieved through incubating slides in citrate buffer at 100 °C for 20 minutes. Slides were blocked in phosphate-buffered saline with 0.05% Tween-20 (PBST0.05) and 3% bovine serum albumin (BSA) for 1 hour at RT. Primary antibodies against CA19-9 (clone 9L426, US Biological Life Sciences) and TRA-1-60 (Novus Biologicals) were labeled for immunofluorescent staining with Sulfo-Cyanine5 NHS ester and Sulfo-Cyanine3 NHS ester respectively. After dialysis to remove unreacted conjugate, the antibodies were diluted into the same solution of PBST0.05 with 3% BSA to a final concentration of 10 μg/mL. Slides were incubated overnight with this solution at 4 °C in a humidified chamber.

The following day, the antibody-containing solutions were decanted and the slides were washed twice in PBST0.05 and once in 1X PBS, each time for 3 minutes. The slides were dried via blotting and incubated with DAPI at 10 μg/mL in 1X PBS for 15 minutes at RT. Two five-minute washes were performed in 1X PBS, and then slides were cover-slipped and scanned using a fluorescent microscope (AxioScan.Z1, Zeiss, Oberkochen, Germany). The microscope collected 3 images at each field-of-view, each image corresponding to the emission maxima of Hoechst 33258, Cy3, and Cy5. We next quenched the fluorescence using 6% H_2_O_2_ in 250 mM sodium bicarbonate (pH 9.5-10) and performed another round of immunofluorescence using two different antibodies. The subsequent incubations and scanning steps were as described above.

Prior to the second round of detection with the TRA-1-60 antibody, we treated the slides with sialidase to remove terminal sialic acids. The slides were incubated with a 1:200 dilution (from a 50,000 U/mL stock) of α2-3,6,8 Neuraminidase in 5 mM CaCl_2_, 50mM pH 5.5 sodium acetate overnight at 37 °C. The subsequent incubations and scanning steps were as described above. The hematoxylin and eosin (H&E) staining followed a standard protocol.

### 2.3 Image processing and analysis

All image data were quantified using SignalFinder. For each image, SignalFinder creates a map of the locations of pixels containing signal and computes various values for the output report, such as the percentage of tissue-containing pixels that have signal. This analysis required major computational resources, as each high-resolution image of the ~1.5 x 2.5 cm tissue comprises about 9 billion pixels at ~3000 pixels/mm resolution and ~500 GB file size. A 1-square-inch image at such resolution would scale to about 4000 square feet if changed to 72 pixels per inch as used for display.

### 2.4 Software

We developed the SignalFinder software using MATLAB, Java, and C++. We used Microsoft Excel and MATLAB for analyzing numerical output; GraphPad Prism, the R language, and Microsoft PowerPoint for the preparation of graphs; and Canvas XIV for the preparation of figures

## 3 Results

### 3.1 Automated signal quantification reveals diverse but non-predictive glycan patterns

We analyzed whole-block tumor sections from 22 patients who underwent resection and adjuvant chemotherapy as curative treatment for PDAC (Table 1). We immunostained each specimen for the CA19-9 and STRA biomarkers and detected the biomarkers using multicolor fluorescence imaging (Fig. 1A). We then analyzed the fluorescence images by automated signal identification (23) to produce maps of the signal pixels (Fig. 1B), which we overlaid onto the brightfield images of the stained slides (Fig. 1B). This system enables a visualization of the locations of biomarker staining and of the histomorphologies of the cells producing the biomarkers (Fig. 1B). But more importantly for the investigations of biomarker features associated with outcomes, the method provided automated, objective signal identification and quantification for use in subsequent biomarker analyses. This foundation allowed us to ask whether we could achieve a fully data-guided method of ranking the likelihood of recurrence, that is, a method of classifying the tumors that does not involve user selections of locations or settings.

**Figure 1.**
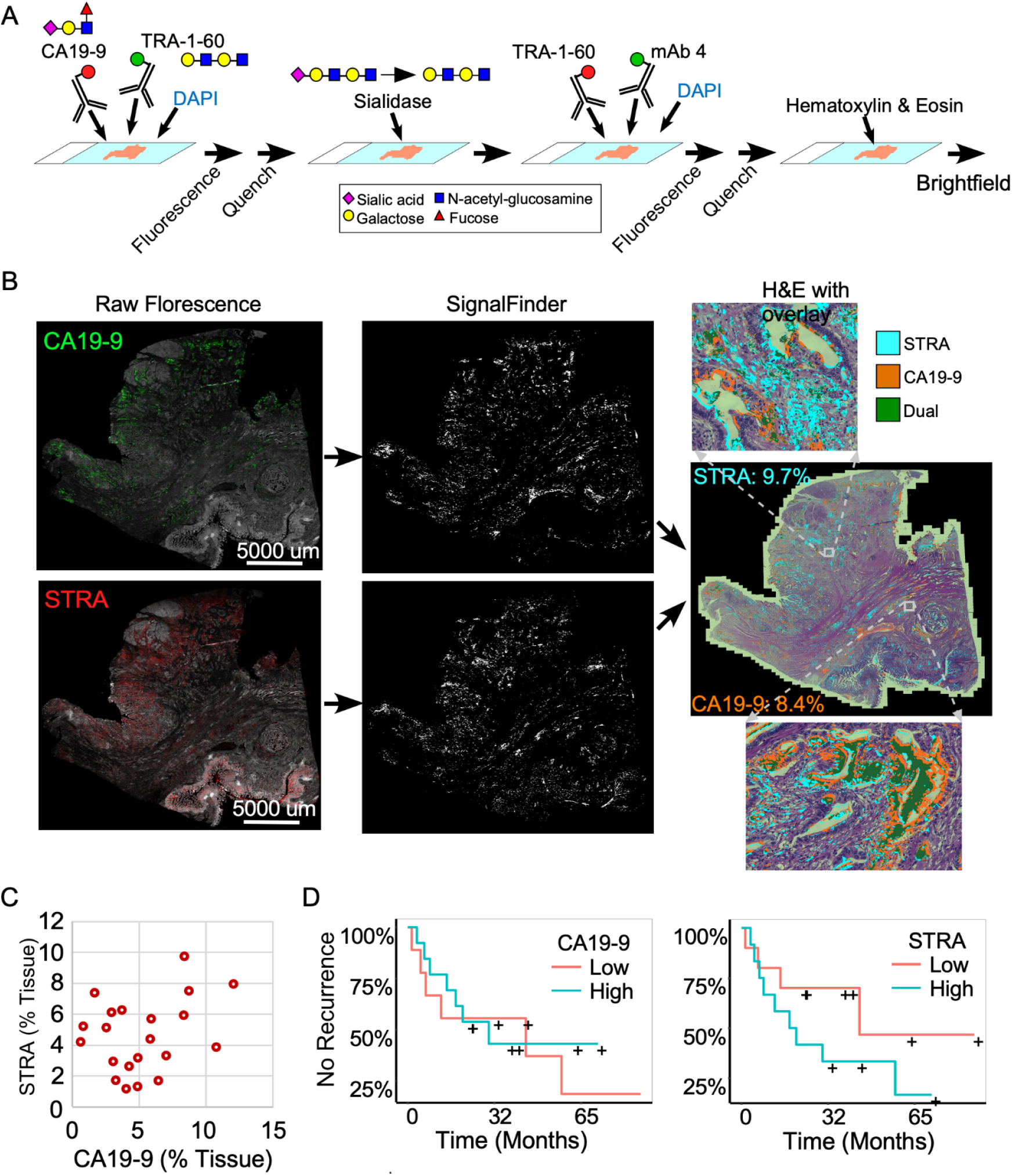
Automated quantification of multiplexed immunofluorescence in whole-block tumor sections. (A) Method of data acquisition. CA19-9 is detected in the first round of staining, and STRA is detected in the second round using the TRA-1-60 antibody post-sialidase. (B) Data processing. The raw fluorescence (left) is processed by SignalFinder to identify signal pixels (middle), which are then assigned colors and overlaid on the H&E brightfield image (right). The percentages are the amount of each signal relative to tissue area. (C) STRA signal amount with respect to CA19-9 amount. Each point is a whole-block tissue specimen. (D) Kaplan-Meier associations of CA19-9 and STRA with time to recurrence. The median value of each defined the cutoff between high and low.

**Table 1.** Cohort information.

| Group | Total | Recurrence | No Recurrence |
| --- | --- | --- | --- |
| Total Samples | 21 | 13 | 8 |
| Avg. Age | 68.2 | 65.5 | 68.3 |
| Percent Male | 42.9% | 38.5% | 50.0% |
| Outlook/Outcome |  |  |  |
| Recurrence < 1 year, N (%) | 7 (33.3) | 7 (33.3) | - |
| Recurrence < 2 years, N (%) | 11 (52.4) | 11 (52.4) | - |
| Recurrence < 3 years, N (%) | 12 (57.1) | 12 (57.1) | - |
| Survival < 1 year, N (%) | 3 (14.3) | 3 (14.3) | - |
| Survival < 2 years, N (%) | 5 (23.8) | 5 (23.8) | - |
| Survival < 3 years, N (%) | 6 (28.6) | 6 (28.6) | - |
| Treatments |  |  |  |
| Gem, N (%) | 14 (66.7) | 12 (92.3) | 2 (25.0) |
| FOLFIRINOX, N (%) | 11 (52.4) | 6 (46.2) | 5 (62.5) |
| Abraxane, N (%) | 8 (38.1) | 8 (61.5) | - |
| FOLFOX, N (%) | 2 (9.5) | 2 (15.4) | - |
| Xeloda, N (%) | 6 (28.6) | 6 (46.2) | 2 (25.0) |
| Olaparib, N (%) | 1 (4.8) | 1 (7.7) | - |
| Radiation, N (%) | 4 (19.0) | 2 (15.4) | 2 (25.0) |

The amounts of each biomarker—as determined by the number of signal pixels identified by SignalFinder normalized to the pixels in the tissue—were highly variable among the 22 subjects and between the biomarkers (Fig. 1C). The subjects had widely varying outcomes, with some exhibiting no evidence of disease after several years and others succumbing to recurrent disease after less than one year (Table 1 and Supplementary Table 1). A group between these extremes had extended survival but struggled with recurrence and continued progression of the disease. The outcomes did not associate with any demographic factors or type of systemic therapy (Table 1). Neither the individual biomarkers (Fig. 1D) nor the combination of the two (not shown) was associated with outcome. An alternative way to quantify the signal using average intensities above a threshold gave different quantifications but also produced no associations with outcome (Supplementary Fig. 1).

### 3.2 Spatial clustering shows divergent types and amounts of glycan clusters

To gain insights into the sources of variability between the specimens, we examined the images of the whole-block sections (Fig. 2). At the microscopic level, we observed extreme diversity within and between tumors in the biomarker locations and amounts. The biomarker staining occurred primarily in epithelial and glandular areas—a potentially useful trait for assessing adenocarcinoma—but it also occurred in varying degrees among both benign and cancerous glands and some non-epithelial area. For example, specimen 17-213 showed CA19-9 staining in ampullary glands and low-grade PanIN and STRA staining in poorly differentiated carcinoma and focally in PanIN, but in another section (16-570), both STRA and CA19-9 were together in varying relative amounts in cancer glands. STRA staining appeared in some small-intestinal mucosa that was at the edge of the section. STRA also appeared alone or together with CA19-9 in cancer glands in a specimen (18-371). We also observed occasional CA19-9 staining of macrophages (16-570a) and red blood cells (18-371a) and STRA staining of stroma (17-213a). Such diversity appeared in the remainder of the specimens (Supplementary Fig. 2). The above observations indicated that simple quantifications of signal amount or intensity across entire sections would include large areas with low or zero biomarker production, as well as signals from both benign and cancerous glands.

**Figure 2.**
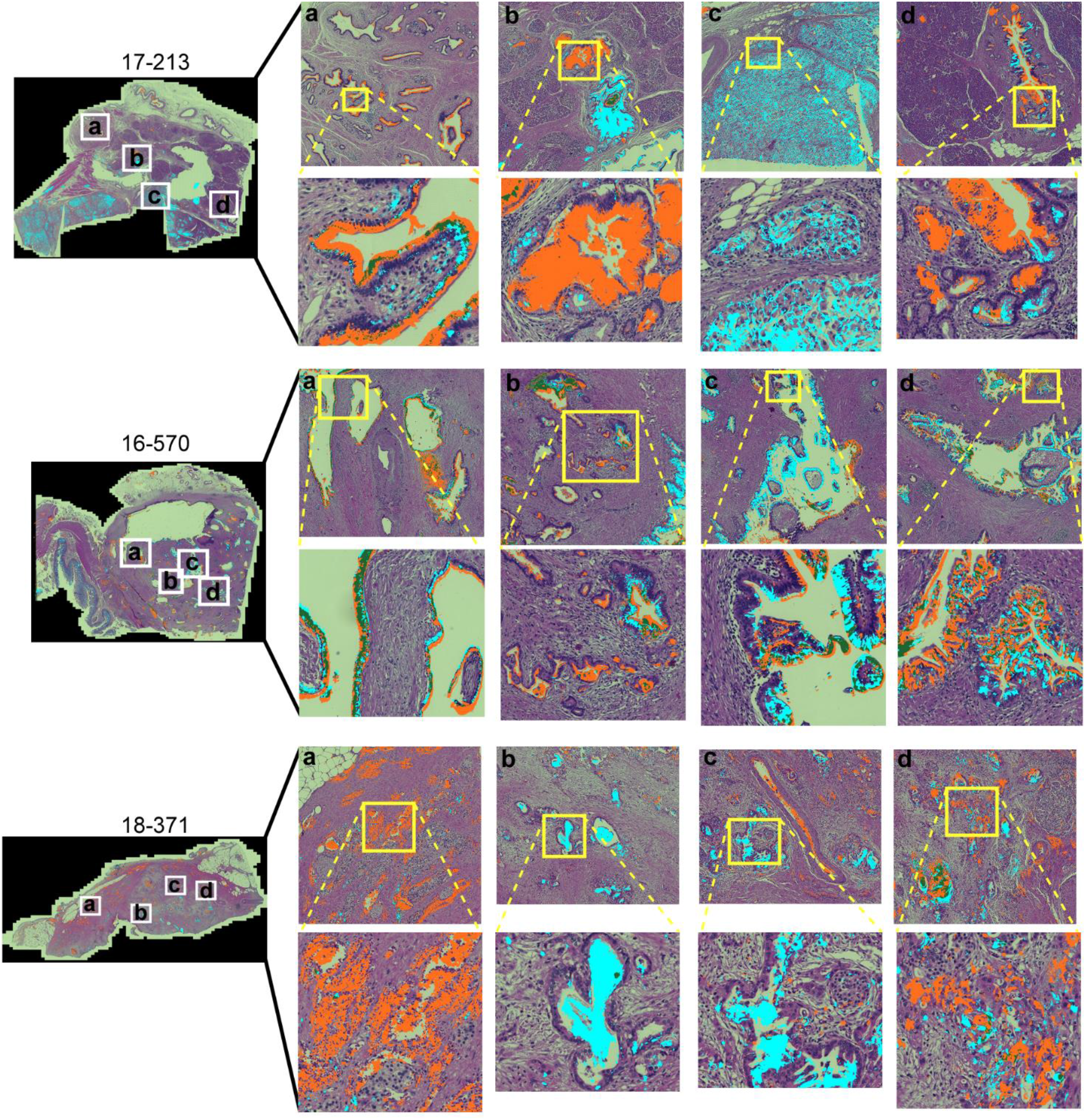
Within-tumor and between-tumor diversity. Medium and tight zoomed regions are shown. The histomorphology shows low grade PanINs (17-213, b and d); benign pancreas (18-371, a); invasive carcinoma (17-213, c; 16-570, a-d; 18-371, b-d); and ampullary glands (17-213, a).

To quantify features that potentially are more specific to tumor assessment, we tallied the signal only where it is part of a cluster of biomarker production (Fig. 3A). Starting with the pixel maps of signal from the SignalFinder output, we produced a sum of signal abundance within a pixel-by-pixel moving box (Fig. 3B). The number of regions with spatial concentrations above various thresholds provided the amount of clustering for each biomarker (Fig. 3B).

**Figure 3.**
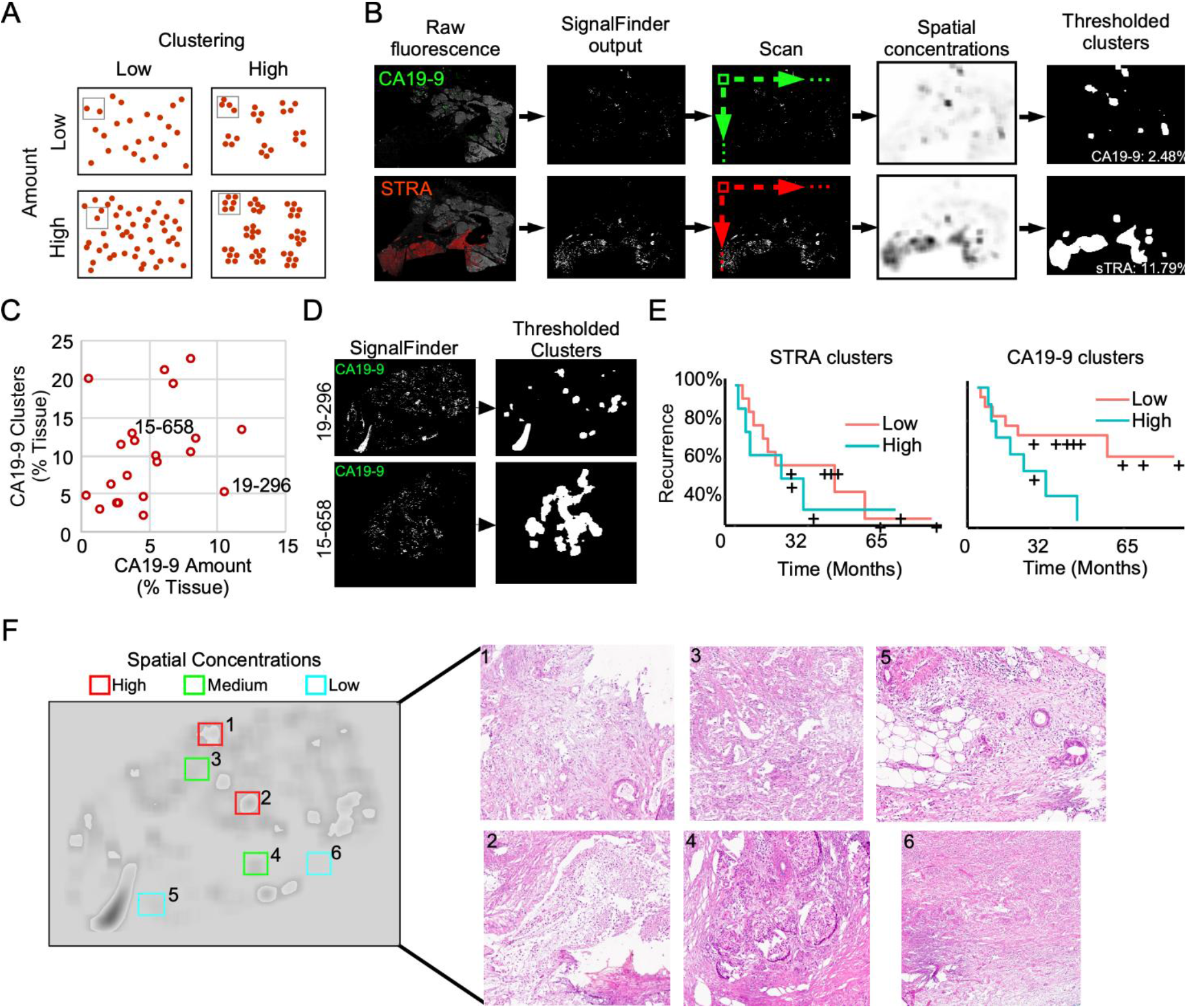
Identification of spatial clusters. (A) Types of relationships between signal amount and clusters. (B) Method of analysis. (C) Relationship between CA19-9 clusters and amounts. Each point represents a specimen. (D) Examples of punctate (19-296) and broad (15-658) clusters. (E) Kaplan-Meier associations of CA19-9 and STRA clusters with time to recurrence. The median value of each defined the cutoff between high and low. (F) Histomorphology in regions defined by spatial concentrations. The left image shows the spatial concentrations of CA19-9. The right panels are corresponding H&E images for the indicated regions, showing invasive carcinoma (panels 1, 3, 5), invasive carcinoma with abundant necrosis (panel 2), benign pancreas (panels 4 and 5), and stroma (panel 6).

We first asked whether this mode of quantification provides distinct information, or alternative whether it simply reflects random clustering that correlates with total signal amount. A view of cluster amounts with respect to signal amounts or intensities showed only weak correlation (Fig. 3C). In some cases, the signals were highly clustered, such as the CA19-9 data in patient 15-658, and in other cases, the signals were more evenly distributed, as with the CA19-9 data in patient 19-296 (Fig. 3D). This finding indicates that the level of clustering is distinct from total biomarker amount and that specimens have diversity between them in the amount of biomarker clustering.

The quantifications of clustered signal, however, did not associate with recurrence to a greater degree than signal amount (Fig. 3E). A survey of the histological features of regions with high or low spatial concentrations gave insights into the tumor characteristics identified by this method of quantification (Fig. 3F). The areas with higher spatial concentrations aligned with greater cellularity, and the areas with very low spatial concentrations were frequently free of epithelia, but areas with both high and low spatial concentration contained various levels of cancer and benign cells. These observations indicated that spatial concentration specifically quantifies the glandular-epithelial biomarker production but does not, by itself, distinguish types of histomorphology.

Given the varied relationship between the glycans (Fig. 2), we next asked whether the combined CA19-9 and STRA provide unique information about the biomarker clusters. We used the maps of the individual biomarker clusters to define the overlap between the glycan-defined clusters and to identify the regions that were exclusively STRA, exclusively CA19-9, or both (Fig. 4A, referred to as STRA-only, CA199-only, and dual). We then separately quantified the amount of each type of cluster to examine whether the overlap was simply a random event correlated with total signal amounts. Some of the tumors just had the STRA-only or the CA199-only clusters (Fig. 4B and Supplementary Fig. 3), and the amounts of clusters containing both markers were not correlated with the sums of the signal amounts (Fig. 4B). For example, subject 15-658 had a low amount of the biomarkers but proportionally high clusters; subject 17-213 largely had the STRA-only clusters, and 18-137 had high amounts and clusters (Fig. 4B). Maps of the three cluster types confirm such various levels of heterogeneity and showed that the different types of clusters can either appear near each other or occupy separate regions (Fig. 4D).

**Figure 4.**
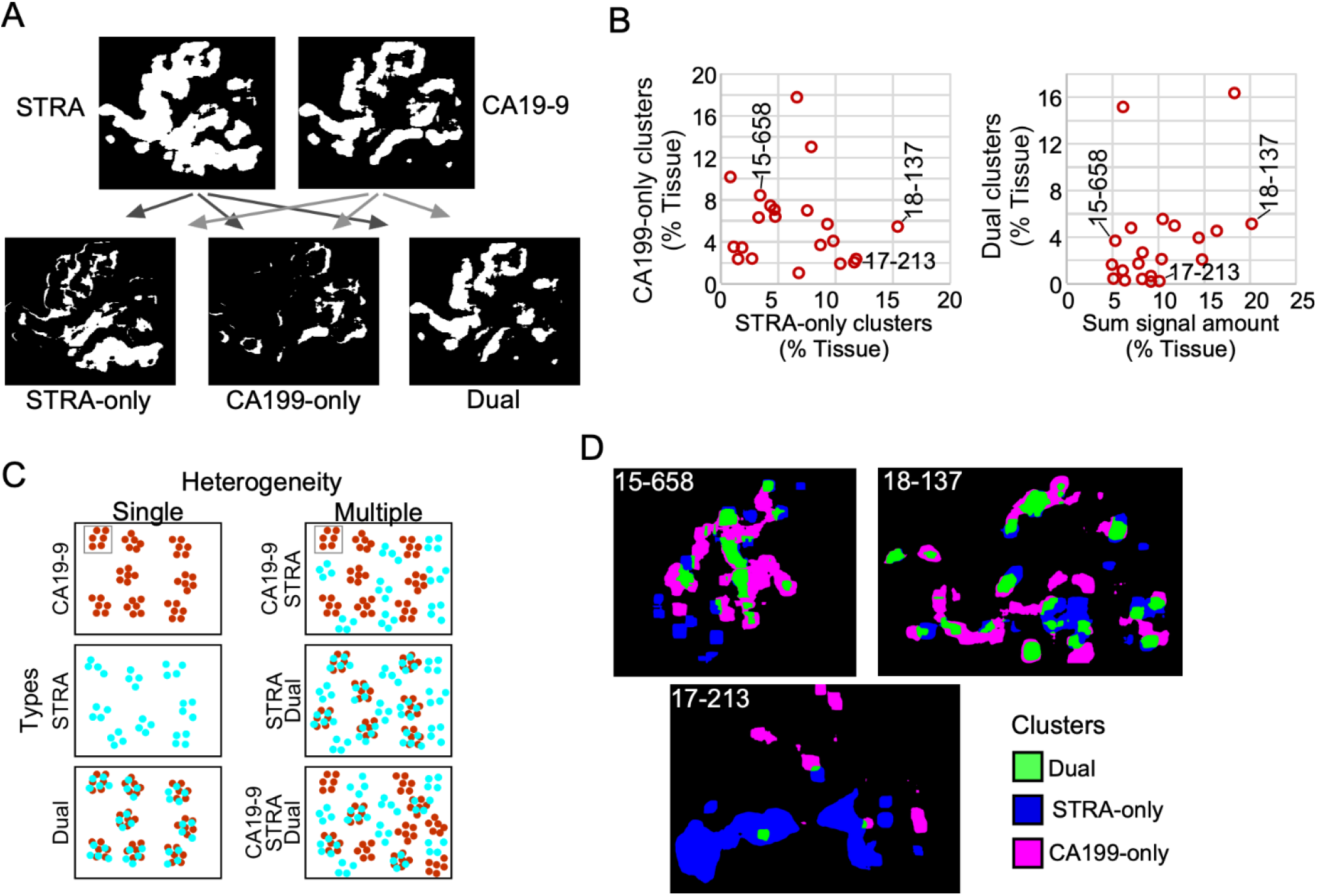
Spatial clusters with distinct biomarker production. (A) Method of identification. The maps of clusters for the two biomarkers were compared to identify the ones high in only one biomarker or in both. (B) Quantification of CA199-only clusters relative to STRA-only clusters (left) and dual clusters relative to sum of signal amount from STRA and CA19-9 (right). (C) Types of cluster compositions in tissue. (D) Visualization of moderate (15-658), high (18-137), and low (17-213) heterogeneity in cluster types.

Given that the dual clusters were not resulting just from a mass-driven overlap of unrelated signals, we concluded that the combined glycans provide distinct information about the biomarker clusters. Furthermore, the specimens vary greatly in the types of clusters they exhibit. Specimens exist that are either relatively homogeneous in the types of clusters they contain or are heterogeneous with various relative proportions of the STRA-only, CA19-9-only, or dual clusters (Fig. 4C).

### 3.3 Heterogeneity in cluster types is associated with outcome

The above findings suggested that a combined evaluation of the distinct cluster types may provide value for tumor assessment. An alignment of the individual biomarker amounts and clusters by subject showed the diversity between subjects (Fig. 5A), both for subjects with and without recurrence of disease after 3 years. It also showed the lack of correspondence between the two methods of quantification, as with subject 19-296, who had a high amount but low clusters of CA19-9. None of the individual biomarker clusters was significantly associated with outcome (Fig. 5B), prompting us to ask whether certain combinations of clusters have stronger associations with outcome than others. To categorize the subjects based on combined biomarker amounts or clusters, we dichotomized the presence or absence of each biomarker using cutoffs determined from receiver-operator-characteristic analysis (Supplementary Table 2). The resulting values illustrated the prominence of only one cluster type in certain tumors and two or more in others (Fig. 5C). Furthermore, it was evident that the co-occurrence of multiple cluster types was a dominant feature of the recurrent specimens. Accordingly, the occurrence of more than one type of cluster was significantly greater in the cases than in controls (11/13 cases vs. 2/8 controls, c^2^ = 7.5, p = 0.006). The improved classification accuracy resulted from the identification of heterogeneity in biomarker clusters in cases where biomarker amounts were not heterogeneous. For example, subject 18-371 had high biomarker amounts only of CA19-9 but high clusters of all three types (Fig. 5C). An equivalent approach applied to the marker amounts resulted in no significant difference (Fig. 5D).

**Figure 5.**
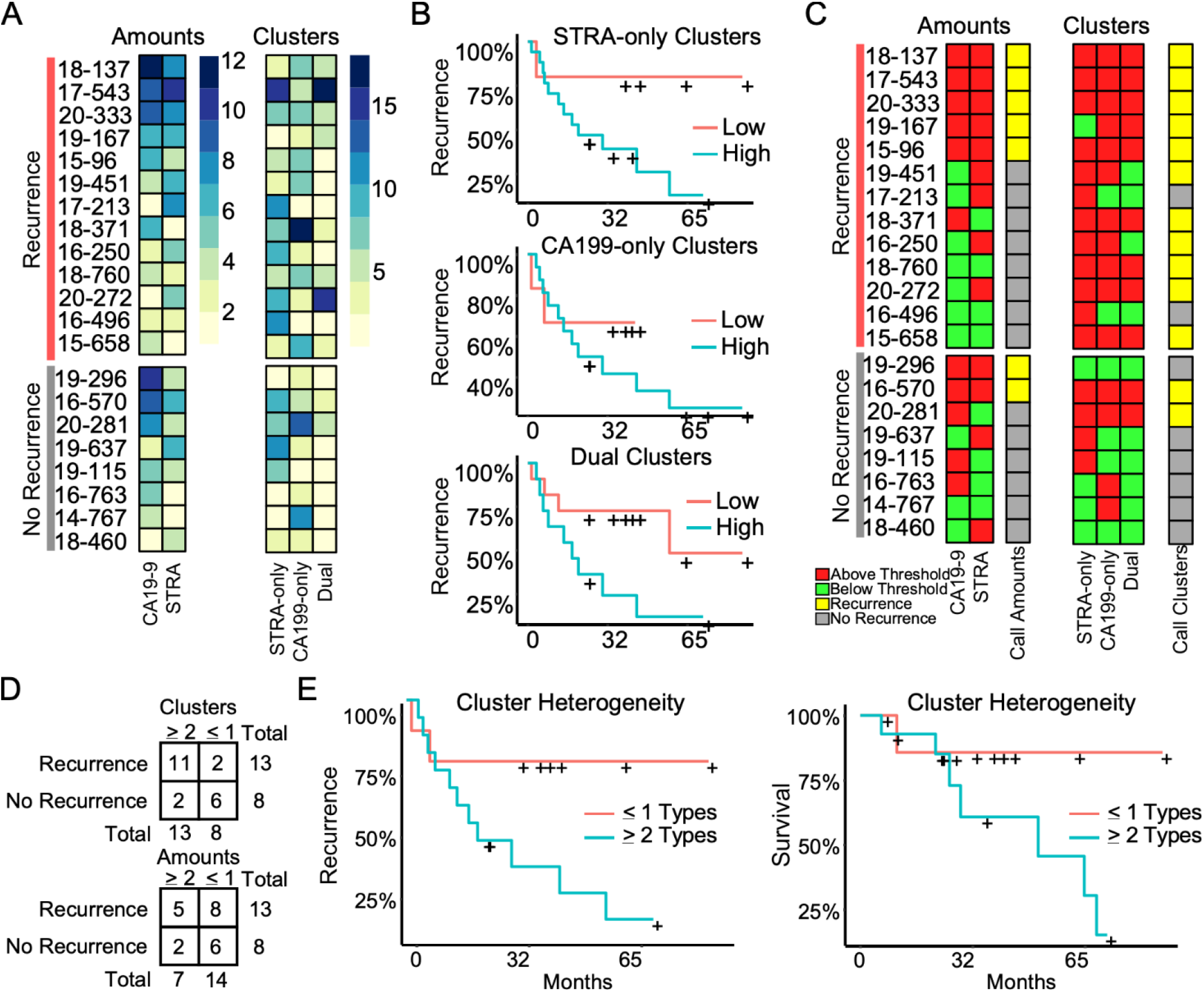
Associations with outcome. (A) Alignments of signal amounts and clusters of each biomarker and cluster type. (B) Associations of single cluster types. (C) Thresholded values and assignment of case status based on >1 biomarker or cluster type. (D) Assignment accuracy using clusters or amounts. (E) Recurrence and survival comparison of subjects with ≥2 cluster types to patients with ≤1 cluster type.

We further tested this relationship using Cox Proportional Hazards models for continuous analysis of survival with respect to spatial cluster abundance (Fig. 5E). Because p values are unstable for survival analysis using small sample sizes, we utilized explained randomness in proportional hazard models of Xu and O’Quigley 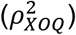 (27). A model using all the spatial cluster abundances indicates improved explanation of survival probability 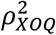 0.52 compared to 0.10, 0.04, and 0.38 for STRA, CA19-9 and Dual, respectively). The explained survival probability surpasses that of the total biomarker amounts 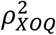 0.16) and average intensities of 90^th^ percentile 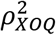 0.10) as well as the individual spatial clusters (non-exclusive 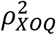 0.28). Because the quantifications of clusters and amounts start from the same SignalFinder output, this result indicates that biomarker clusters are more informative for outcomes than biomarker amounts and that heterogeneity in biomarker clusters is an indicator of rapid progression in PDAC.

We tested the reproducibility of these results for 5 new tissue specimens that were cut from the same blocks as used in the above analysis but separated in the block by 10-20 mm. The quantifications of clusters correlated positively between the first and second runs, and after applying thresholds to classify each case, the classifications matched exactly (Supplementary Figure 4). We concluded that for these sections, the types of biomarker clusters are consistent between different sections of a tumor block within moderate variation.

### 3.4 Regional cluster heterogeneity aligns with pathology-confirmed cancer in recurrent tumors

The association of cluster heterogeneity with recurrence suggests that the co-occurrence of divergent clusters identifies a subset of cancer cells. We therefore investigated whether areas with biomarker cluster heterogeneity are more associated with the presence of cancer cells than areas without cluster heterogeneity. We performed intra-tumoral comparisons of histomorphology in 4 specimens with heterogeneity in clusters but not in amounts and in 1 specimen with heterogeneity in both clusters and amounts (Fig. 6). For each, we identified regions containing more than one cluster type and regions containing just one cluster type within regions-of-interest (~3 mm) three times the size of the region used in calculating spatial concentrations (~1 mm, Fig. 3B). We then identified by surgical pathology review the regions containing cancer glands and compared the overlap with the clusters. At the macroscopic level, the areas with heterogeneous clusters corresponded well with the locations identified by surgical pathology as containing cancer cells (Fig. 6A). In three subjects (16-250, 15-658, and 18-371), the identifications were equivalent, but in the others, cluster heterogeneity occurred in some regions not identified by surgical pathology. In all, cluster heterogeneity identified areas containing cancer cells in 28 out of 32 (88%) instances. The matched regions-of-interest taken from areas containing only 1 cluster type overlapped with cancer-cell-containing regions in 2 of 17 (11.7%) instances. This result indicates that heterogeneity in biomarker clusters preferentially identifies regions containing cancer cells, in contrast with the individual biomarkers.

**Figure 6.**
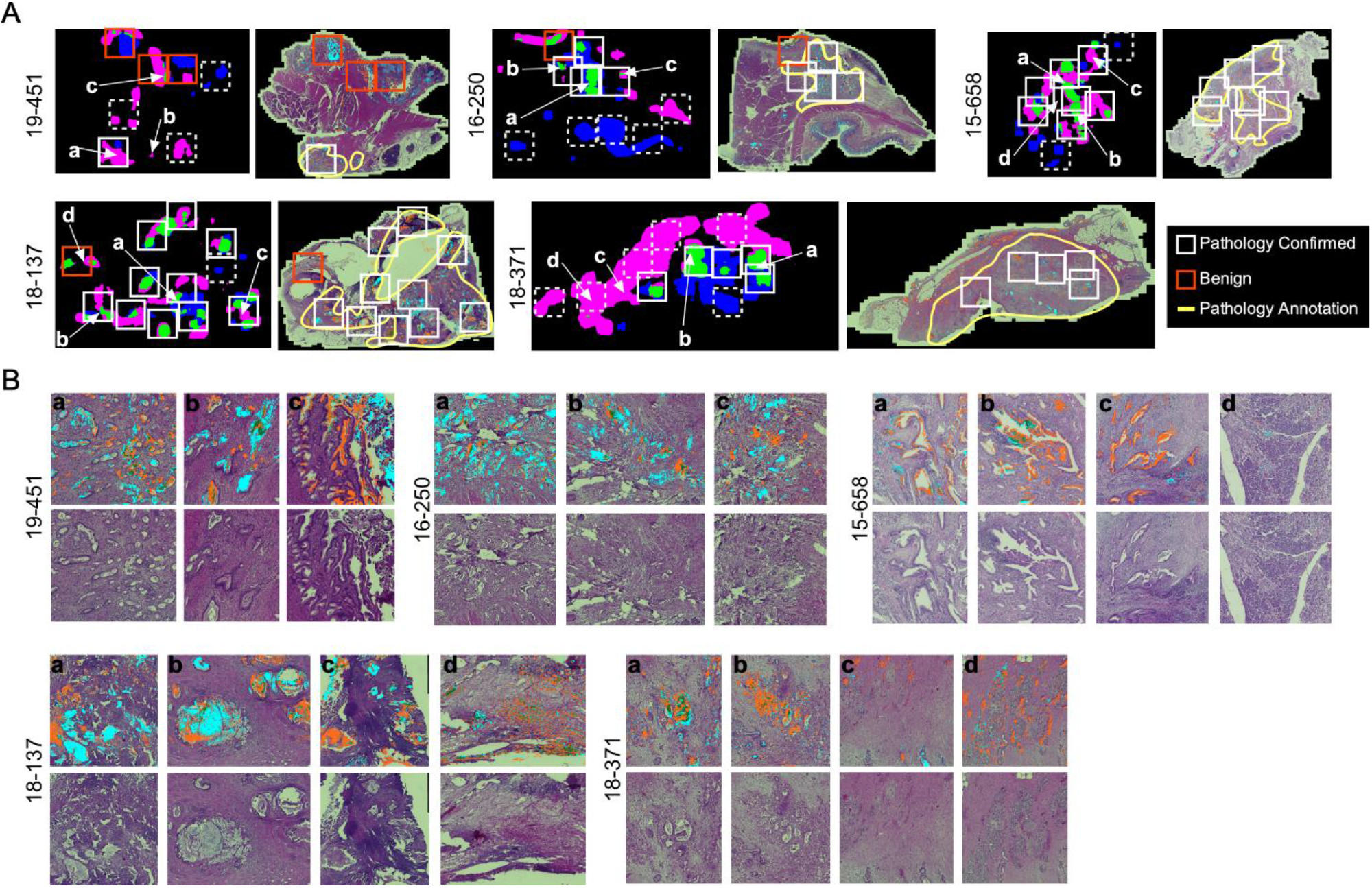
Comparison to histopathology. (A) General localization of cancer-container areas. The white boxes indicate regions of tissue with >1 cluster type within areas of pathology-confirmed cancer; the red boxes indicate similar areas outside of pathology-confirmed cancer; and the dashed boxes indicate regions with <2 cluster types. The yellow lines indicate regions as broadly containing cancer cells. (B) Microscopic-level comparison of regions identified by cluster heterogeneity and histopathological review.

Given that cluster heterogeneity associated with recurrence, we further examined the histological differences between the recurrent and non-recurrent specimens. By surgical pathology, no systematic differences were evident between the patients with recurrence and without recurrence, nor between the patients with cluster heterogeneity and without cluster heterogeneity (Table 2). We concluded therefore that heterogeneity in biomarker clusters provides information about cancer-containing regions that is not discernable from histopathological review, in particular as a quantifiable feature of tumors with high likelihood of recurrence.

**Table 2.** Comparison with surgical pathology. Outcome is the case/control status from Table 1. *Cluster types refers to the number of different types of clusters (from Figure 5D). Shaded cells are those with <2 cluster types.

| ID | Outcome | Cluster types* | Site | Grade | Margins | Invasion | Lymph nodes | Peri-neural | T | N | M |
| --- | --- | --- | --- | --- | --- | --- | --- | --- | --- | --- | --- |
| 17213 | Case | 1 | Head | Poorly Differentiated | Positive | Positive | Positive | ND | 3 | 1 | 1 |
| 20272 | Case | 3 | Tail | Moderately Differentiated | Negative | Positive | Negative | Positive | 2 | 0 | X |
| 18760 | Case | 3 | Head | Moderately Differentiated | Negative | Positive | Positive | Positive | 2 | 2 | X |
| 16496 | Case | 1 | Head | Moderately Differentiated | Negative | Positive | Positive | Positive | 3 | 1 | X |
| 18137 | Case | 3 | Tail | Moderately Differentiated | Positive | Positive | Negative | Positive | 3 | 0 | X |
| 19167 | Case | 2 | Tail | Moderately Differentiated | Positive | Positive | Positive | Positive | 2 | 1 | X |
| 18371 | Case | 3 | Tail | Moderately Differentiated | Negative | Negative | Negative | Positive | 1 | 0 | X |
| 1596 | Case | 3 | Tail | Moderately Differentiated | Negative | Positive | Negative | Positive | 2 | 0 | X |
| 20333 | Case | 3 | Head | Moderately Differentiated | Negative | Positive | Negative | Positive | 2 | 0 | X |
| 19451 | Case | 2 | Head | Moderately Differentiated | Negative | Positive | Positive | Positive | 3 | 2 | X |
| 16250 | Case | 2 | Head | Moderately Differentiated | Positive | Positive | Negative | Negative | 3 | 1 | X |
| 17543 | Case | 3 | Head | Moderately Differentiated | Negative | Positive | Positive | Positive | 3 | 1 | X |
| 15658 | Case | 3 | Head | Moderately Differentiated | Positive | Positive | Positive | Negative | 3 | 1 | X |
| 20281 | Control | 3 | Head | Well Differentiated | Negative | Positive | Positive | Positive | 2 | 1 | X |
| 19637 | Control | 1 | Head | Poorly Differentiated | Negative | Negative | Positive | Positive | 2 | 1 | X |
| 19296 | Control | 0 | Tail | Moderately Differentiated | Negative | Positive | Positive | Positive | 2 | 1 | X |
| 19115 | Control | 1 | Head | Moderately Differentiated | Negative | Negative | Negative | Positive | 2 | 0 | X |
| 18460 | Control | 0 | Tail | Moderately Differentiated | Negative | Negative | Negative | Negative | 2 | 0 | X |
| 16763 | Control | 1 | Tail | Moderately Differentiated | Negative | Positive | Positive | Negative | 3 | 0 | X |
| 14767 | Control | 1 | Head | Moderately Differentiated | Negative | Positive | Positive | Positive | 3 | 1 | X |
| 16570 | Control | 3 | Head | Moderately Differentiated | Negative | Positive | Positive | Positive | 3 | 1 | X |
| 20282 | Unknown | 2 | Head | Moderately Differentiated | Positive | Positive | Positive | Positive | 1 | 2 | X |

## 4 Discussion

The keys to uncovering accurate predictors of outcome are markers of the individual, relevant subpopulations and a method to quantify the relationships within a tumor that are informative of tumor behavior. Here we demonstrate that two glycan biomarkers – CA19-9 and STRA – are predictive of outcome when detected in a particular intra-tumoral relationship. Using a new approach to analyze and quantify biomarker patterns in multiplexed immunofluorescence data, we found that simple quantifications of biomarker amounts were not sufficiently informative of recurrence. But by quantifying the spatial clusters of the combined biomarkers, we found that the co-occurrence of more than one type of biomarker cluster within a tumor associated with recurrence, in contrast to the individual biomarker clusters or glycan levels. These results demonstrate the value of the CA19-9 and STRA glycans for assessing tumors, and they present a new method for quantifying features within tumors that are indicative of tumor behavior.

A plausible interpretation of the relationship found here is that CA19-9 and STRA in combination are markers of clonal heterogeneity of cancer cells. This interpretation concords with previous findings that the two glycans could identify both a stem-like founder population (marked by STRA) and another more-differentiated population (marked by CA19-9) (24). This link also concords with previous studies demonstrating the aggressive nature of clonal heterogeneity in tumors. Heterogeneity in cancer cells, which potentially reflects the plasticity and outgrowth of stem-like cancer cells (12,25), likely provides cells greater ability to adapt, produce diverse subpopulations, and survive. Consistent with these concepts, mouse pancreatic cancer cells gaining plasticity through GATA6 loss increased their chemoresistance and ability to escape immune elimination (26), and tumors able to switch subtypes through the expression of the GLI2 transcription factor have shorter survival and higher tumor growth rates (27). Cancer progression could be further aided not just by heterogeneity in the cancer cells, but also in the tissue microenvironment as defined by fibroblast differentiation, immune activation, and cancer-cell markers (28).

Additional markers could potentially provide new biological information or predictive capability, such as markers of cancer subtypes. For example, GATA6 expression marks the classical type (29,30) and is suppressed in the basal type (26), and a TP63 isoform called deltaN-P63 is suppressed in the classical PDAC transcriptional program (31,32) and could mark a subset of basal-like cancer cells. Neutrophil infiltration (33), epigenetics traits, and metabolites (9,26,34,35) also mark subtypes of cancer. Greater value could be derived through linking with spatial transcriptomics methods. Such methods are powerful for uncovering biological functions and biomarkers (36,37) and, if coupled with the precision and resolution of immunofluorescence, would provide more information about the cells that produce each biomarker.

The current study has several limitations and areas for further development. In the first place, the sample set is small. Future validation studies should be designed with larger cohorts and blinded outcomes that are revealed after analyses are complete. A second limitation and goal for future research is that we did not account for microenvironment. Future studies should include immune cell and fibroblast markers that potentially mark the sub-TMEs (28) or subtypes of stroma found by gene-expression profiling (10). Third, we did not perform a full exploration of the algorithms and parameter space for defining spatial clusters and heterogeneity, owing to the scale of such an undertaking. Here we demonstrated one method that showed the value of the approach, and several of the parameters should be further optimized. In addition, other software programs potentially could examine these questions from other angles, especially those that offer segmentation for the counting of cells. Certain morphological features of PDAC cells associated with subtypes and outcomes (38), suggesting that segmentation methods to identify such feature could augment biomarker discovery algorithms. Many other spatial clustering approaches are available to examine aggregation of particular cell types (39). Such approaches would provide additional layers of interpretation.

Another goal for future work is to broaden the application beyond patients who had resection specimens available. Samples that are available before surgery include biopsy material and peripheral blood. Both types of samples could be amenable to the methods demonstrated here. Biopsy specimens likely would contain the heterogeneous biomarker types within one or more samples, considering that the distinct types of biomarker clusters appeared together within common regions of the tissue (Fig. 6). Blood specimens also could contain the information of cluster heterogeneity, as PDAC cells secrete multiple types of biomarkers into the blood, such as genomic DNA, extracellular vesicles, proteins, metabolites, and others. Glycans in particular have value as markers of cell type that appear both on cell surfaces and in secretions (40,41). In future research, the tissue analysis method demonstrated here could identify the biomarker combinations that are useful for patient evaluation and that could constitute clinical assays using either biopsy or blood specimens. If validated, the method and findings presented here could help to stratify patients by likelihood of recurrence. The tumors in such subtypes could be analyzed for differential responses to the great range in drug options now available or used in the development of patient-derived organoids to identify effective treatments through high-throughput screening (12,42)

## 6 Author Contributions

Performed experiments: L.W. and S.B.; analyzed data: L.W., S.B., Z.K., C.G., C.S., and B.H.; prepared figures and tables: L.W., S.B. and B.H.; Prepared text: B.H.; Contributed resources: P.A.; reviewed paper: all authors.

## 8 Acknowledgments

We thank Dr. Daniel Frobish and Anna Repesh (Grand Valley State University) for assistance in the statistical analysis of associations with outcome.

## 1 Data Availability Statement

The original contributions presented in the study are included in the article/supplementary material. The raw image data the conclusions of this article are available upon request.

