## Supplementary Information for "Heterogeneity of Glycan Biomarker Clusters as an Indicator of Recurrence in Pancreatic Cancer"

### 1 Supplementary Tables

#### 1.1 Supplementary Table 1. Patient Data

| ID | Gender | Age | Treatments | Group | Outcome |
| --- | --- | --- | --- | --- | --- |
| 15-96 | M | 74 | Gemcitabine/abraxane then FOLFOX | Case | Recurrence at 1.5 years; OS 2.5 years |
| 16-496 | M | 73 | Single agent Gemcitabine then  Abraxane | Case | OS < 1 year |
| 17-213 | M | 66 | Gemcitabine and Abraxane | Case | OS < 1 year |
| 18-137 | F | 72 | Gemcitabine and Xeloda; stopped Xeloda, initiated Gemcitabine and Abraxane; FOLFOX | Case | Recurrence < 1 year; OS ~2 years |
| 18-371 | F | 55 | Gemcitabine and Capecitabine; FOLFIRINOX; Gemcitabine and Abraxane | Case | Recurrence and progression < 1 year |
| 18-760 | F | 63 | Capecitabine/radiation then FOLFIRINOX then Gemcitibine/Abraxane | Case | Recurrence at 1.5 years; mets at 2 years |
| 19-167 | F | 41 | FOLFIRINOX then Gemcitibine/Abraxane | Case | Progression at < 1 year; OS < 2 years |
| 20-272 | F | 80 | Gemcitabine and Xeloda | Case | OS < 1 year |
| 15-658 | M | 39 | FOLFIRINOX then Gemcitabine; Gemcitabine/nap-paclitaxel plus trial drug | Case | Recurrence 3.5 years after surgery; OS 5.5 years |
| 16-250 | F | 66 | Capecitabine/radiation then Gemcitabine | Case | Recurrence at 5 years; OS 6 years |
| 17-543 | F | 80 | Gemcitabine; added Cisplatin; radiation with oral capecitabine; Olaparib | Case | Recurrence at ~2 years; OS 4.5 years |
| 19-451 | F | 77 | FOLFIRINOX then Gemcitabine/Abraxane | Case | Recurrence at 1 year; progression at 2 years; stable at 3 years |
| 20-333 | M | 62 | FOLFIRINOX | Case | Recurrence at ~ 2 years |
| 14-767 | M | 73 | Gemcitabine | Control | NED at 7.5 years |
| 16-570 | F | 65 | Capecitabine/radiation then Gemcitabine | Control | NED at 6 years |
| 16-763 | F | 64 | Gemcitabine | Control | NED at 6 years |
| 18-460 | M | 69 | FOLFIRINOX then radiation/Xeloda | Control | NED at almost 4 years |
| 19-115 | F | 70 | FOLFIRINOX | Control | NED at 3 years |
| 19-296 | M | 65 | FOLFIRINOX | Control | NED at 3 years |
| 19-637 | F | 72 | FOLFIRINOX | Control | NED at almost 3 years |
| 20-281 | M | 68 | FOLFIRINOX | Control | NED at almost 2 years |
| 20-282 | F | 66 | FOLFIRINOX | NA | Insufficient follow up |

#### 1.2 Supplementary Table 2. Receiver-operator characteristic analysis to determine cutoffs

See separate Excel file.

### 2 Supplementary Figures

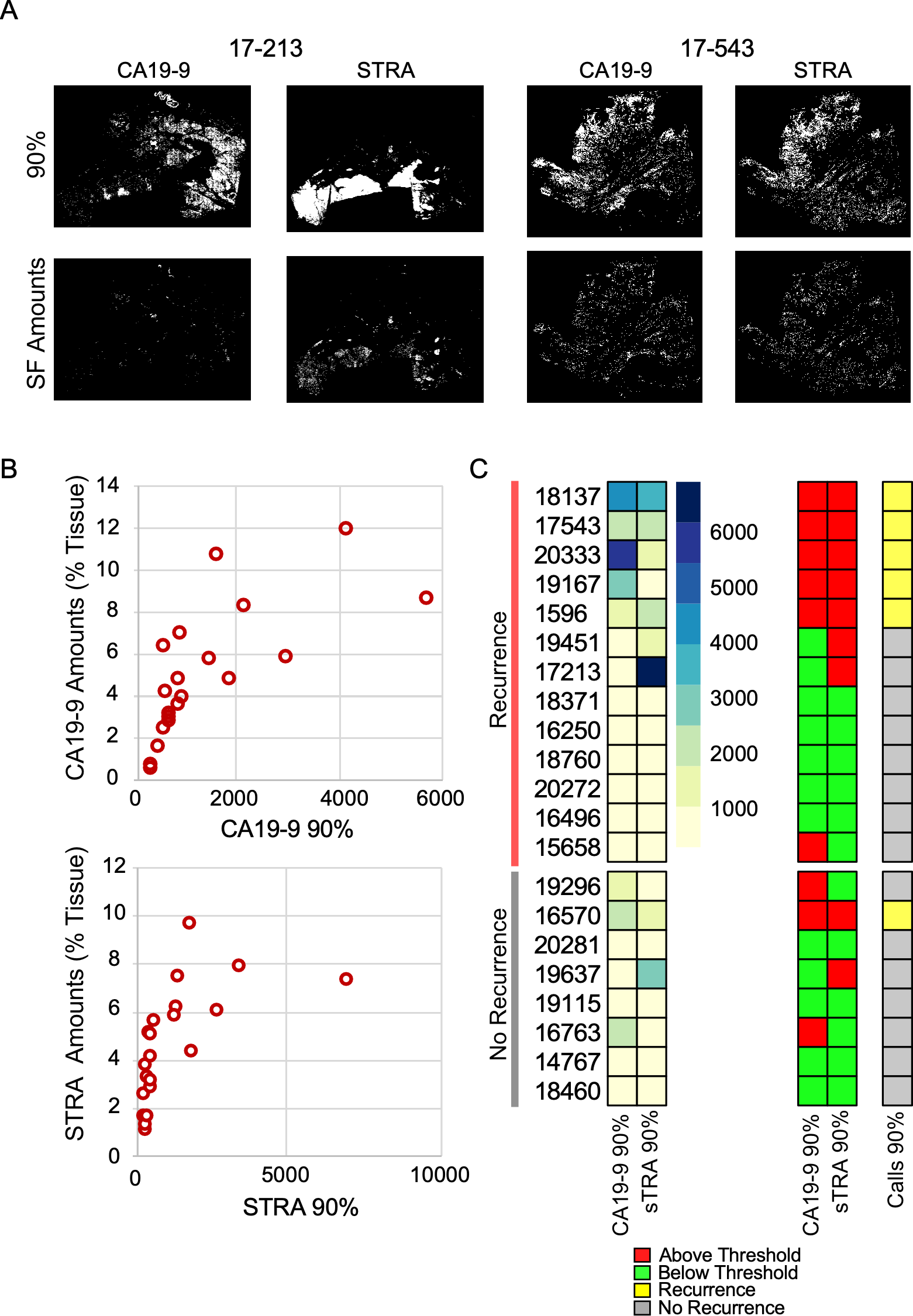

**Supplementary Figure 1.** **Intensity-Based Thresholding.** (A) Comparison of detected signal locations. The top panels show the top 10% of pixels based on intensity, and the bottom panels show the pixels found by SignalFinder. The intensity-based threshold was less selective. (B) Correlation of the two methods of quantification, showing general correlation at lower intensities but much less at higher intensities. (C) Classification using intensity-based thresholds, based on the same system as described in the main text. The accuracy was not statistically significant.

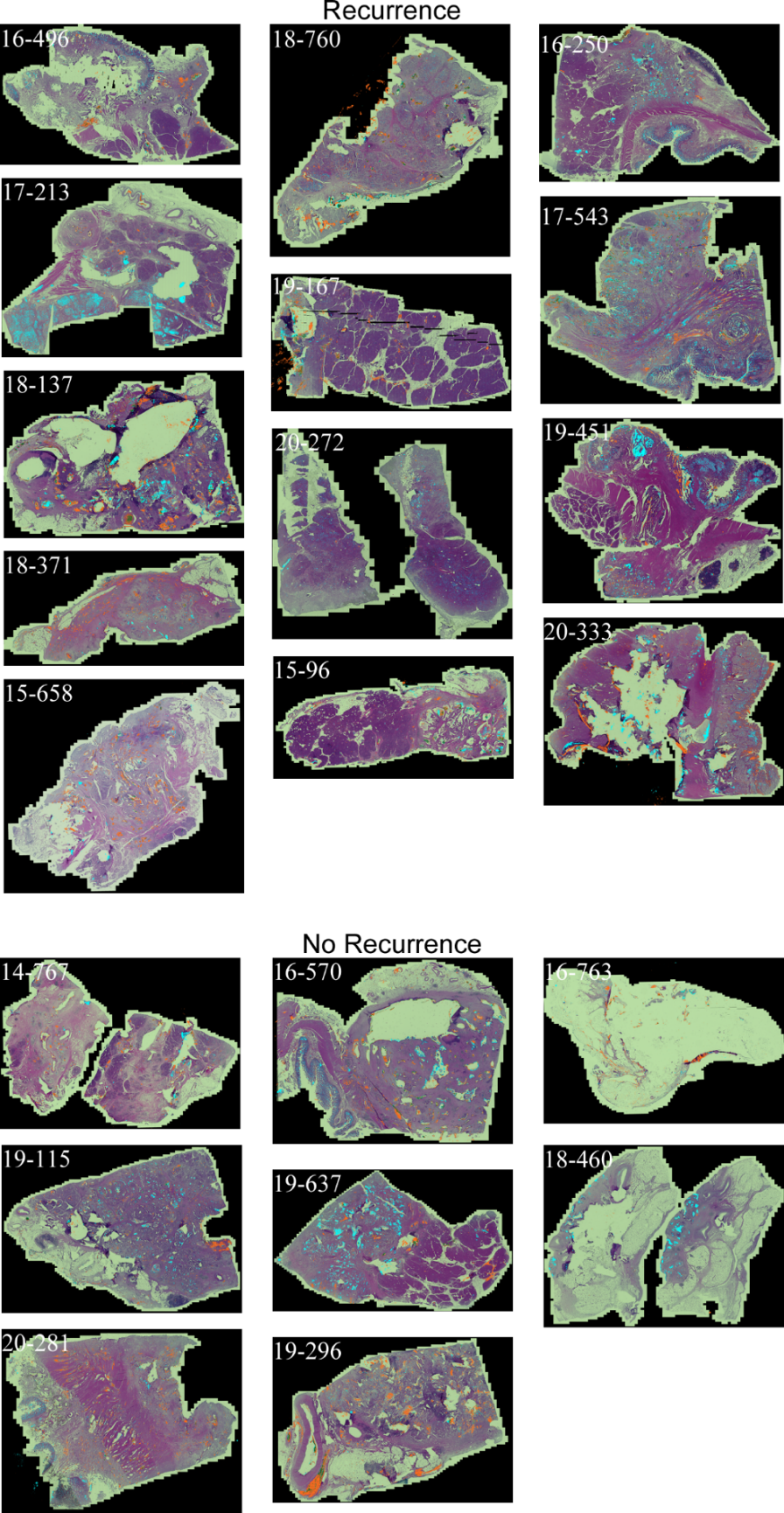

**Supplementary Figure 2.** **Whole-block tumor specimens. The** images show the H&E-stained tissue overlaid with the SignalFinder-detected signals from STRA (cyan) and CA19-9 (orange).

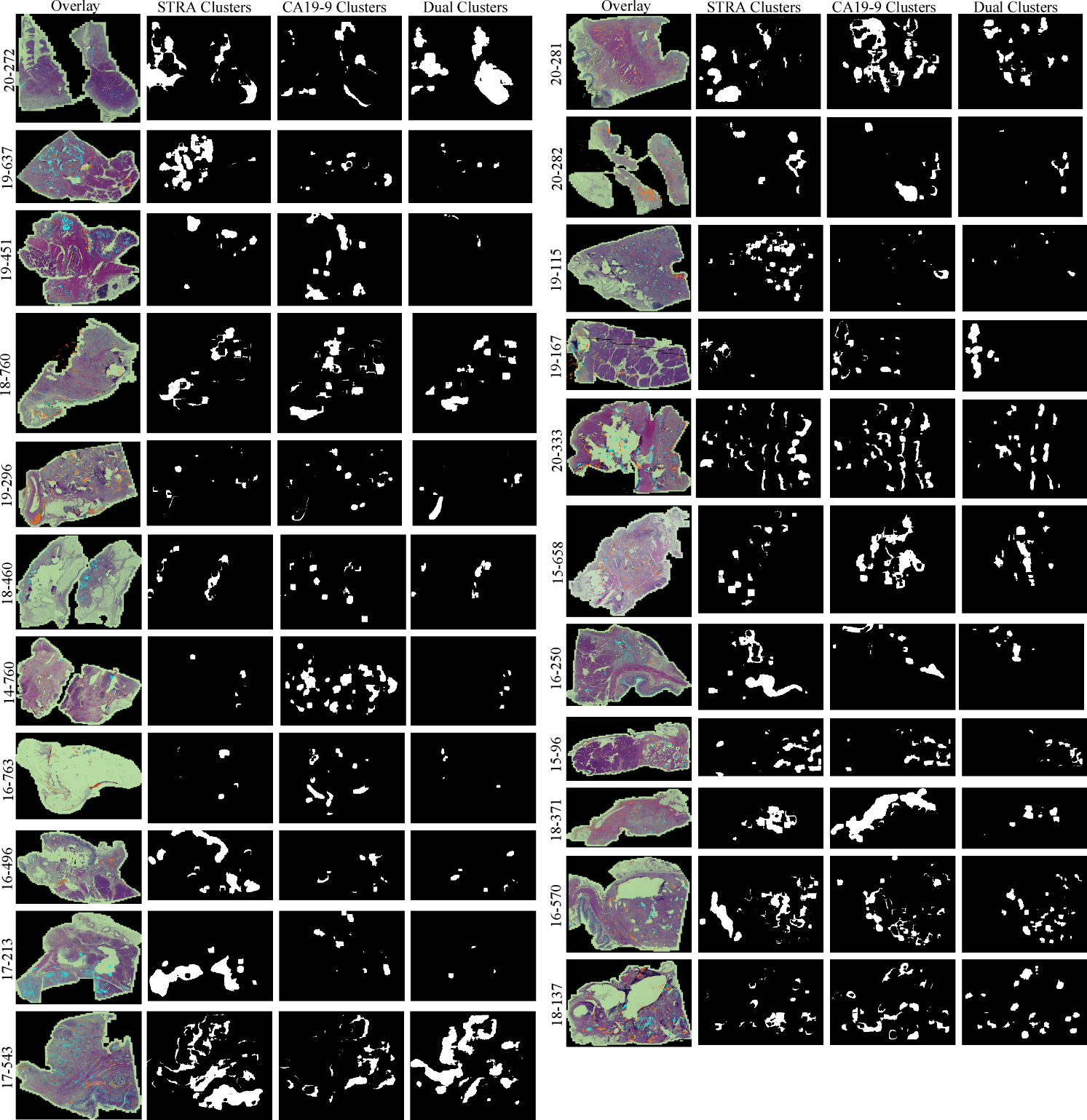

**Supplementary Figure 3.** **Cluster maps.** Each row shows the signal-overlaid H&E image followed by the maps of the STRA-only, CA199-only, and dual clusters.

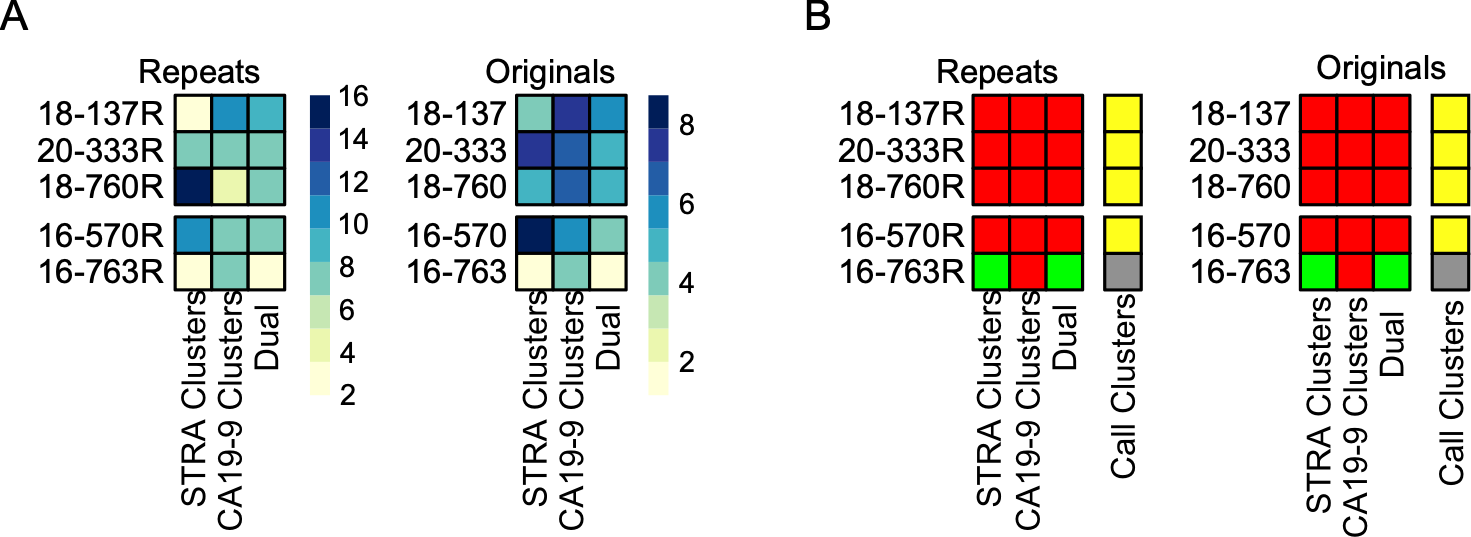
g,. n

**Supplementary Figure 4.** **Test of consistency between sections.** (A) The matrix shows the quantifications of the three types of clusters in the original sections and the repeat sections, taken 10-20 mm removed from the original sections. (B) Thresholded data using the same thresholds and classification rule for both sets.
